# Impact of HIV-1 infection on the microglia transcriptome: induction of microRNA-23b leads to G3BP2-mediated stress granules assembly disruption

**DOI:** 10.64898/2026.10.28.762254

**Authors:** Robert Lodge, William Branton, Frank Kirchhoff, Christopher Power, Éric A. Cohen

## Abstract

Microglia act as HIV reservoirs in the brain and contribute to the development of HIV-associated neurocognitive disorders despite antiretroviral therapy. We examined the transcriptome of primary microglia infected *ex vivo* with HIV-1. Analyses reveal a significant modulation of DNA damage/cell cycle-, cellular stress- and non-membrane bounded organelle assembly-related gene expression, and a broad effect on the microRNA profile. MicroRNA-23b is identified among upregulated microRNAs in HIV-1-infected microglia, and downregulated *g3bp2* mRNA, validated as its target. G3BP2 is a core component of stress granules (SGs), which serve as antiviral and proinflammatory platforms. MicroRNA-23b mimics reduced both G3BP2 expression and SG assembly in stress-induced monocyte-derived microglial cells (MDMi), and few SGs aggregate in infected MDMi. Inhibition of MicroRNA-23b partially re-established SG formation and reduced HIV-1 replication. Thus, microRNA-23b induction contributes to HIV-1 strategies aimed at counteracting SG assembly in infected microglia, a host response that restricts HIV-1 replication and ensuing neuropathogenesis.

**Highlights:**

- Cellular stress response genes are modulated in HIV-infected microglia
- HIV-1 upregulates miRNA-23b, which targets stress granule (SG) factor G3BP2
- MiRNA-23b upregulation is dependent on HIV-1 glycoproteins expression
- Inhibiting miRNA-23b partially restores SGs and reduces HIV-1 replication

## Introduction

Microglia are resident and long-lived innate immune cells of the central nervous system (CNS) that are important for immune surveillance and the homeostatic functions of the CNS^1^. Unlike monocyte-derived macrophages, microglia originate from yolk-sac progenitors that migrate to and proliferate in the CNS^2,3^. *In vivo* evidence indicates that microglia are among the brain cells infected by human immunodeficiency virus type 1 (HIV-1)^4^ and recent studies revealed that microglia harbor replication-competent HIV-1 in virally-suppressed individuals, thus supporting their role as the major persistent cellular reservoir in the brain ^5–7^. In line with viral analyses showing that CNS cells are predominantly infected with macrophage-tropic HIV-1 (R5-M-tropic) variants that can utilize the CCR5 co-receptor and low surface densities of the CD4 receptor for efficient viral entry and infection^8,9^, several primary microglial cell models express CD4 and CCR5 and are susceptible to R5-M-tropic HIV-1 productive infection *in vitro* ^10^. The long-term impact of infection on microglial cell function and its resulting persistent proinflammatory response is thought to contribute to a wide range of CNS manifestations, collectively termed HIV-associated neurocognitive disorders (HAND), which is still prevalent in up to 20% of anti-retroviral therapy (ART)-treated HIV-infected individuals^11–14^. Understanding the complex HIV-microglia interactions is important to design new strategies to impede viral persistence in the CNS and prevent HAND.

Recent advances in transcriptomic technologies have led to the identification of numerous putative targets and molecules with therapeutic potential, including microRNAs (miRNAs)^15^. MiRNAs are small regulatory noncoding RNAs that act on gene expression at the posttranscriptional level. Following their multistep biogenesis, mature miRNAs bind mostly to the 3’ untranslated region (3’UTR) of target mRNAs, leading to their degradation or translational repression ^16^. Thus, miRNAs act as important regulators of cellular processes, from apoptosis to cell differentiation and activation. Furthermore, they serve as modulators of immune function, inflammation and response to infections ^17^. Hence, they have important roles in viral infections, including HIV-1, and their expression can be modulated by host or viral factors ^17,18^. Our previous analysis of miRNA profiles in HIV-1 infected macrophages revealed that miRNAs-221/222, which are induced by tumor necrosis factor alpha (TNF-α), acted as key effectors of the host antiviral responses by inhibiting viral entry through the downregulation of the CD4 receptor ^19^. Moreover, these studies highlighted a novel mechanism by which HIV-1 counteracts the restriction mediated by the type 1 interferon (IFN-1)-regulated MARCH-1 E3 ubiquitin ligase on HIV-1 infectivity, which involves downregulation of its expression via HIV-1 Vpu-mediated induction of miRNAs-25/93 ^20^. Thus, miRNAs can contribute to the enhancement or inhibition of viral replication depending on whether their target functions as a HIV-1 restriction factor or a host dependency factor (HDF), respectively.

Among the various host responses modulated following virus infection are the assembly of cytoplasmic ribonucleoprotein biocondensates named stress granules (SGs) ^21^. These membrane-less organelles assemble in response to external stresses, such as heat shock, viral infection, or oxidative stress (for instance, following addition of arsenite (As)) ^22^. SGs consist of untranslated mRNAs, translation initiation factors and RNA-binding proteins ^22^. However, their aggregation is initiated by core factors, including Ras-GTPase-activating protein-binding proteins 1/2 (G3BP1/2) ^23^ and Polyadenylate-binding protein – cytoplasmic 1 (PABPC1)^24^. SGs act as part of the antiviral response by amplifying innate immune signaling cascades and sequestering viral RNAs ^21,25^. Viruses counteract the SGs antiviral effect by targeting their formation or dynamics ^25^; for instance, the viral protein nsP3 of Chikungunya virus and other alphaviruses ^26^ and the SARS-CoV-2 virus nucleocapsid protein ^27^ both bind G3BP1 to inhibit SGs. In the case of HIV-1, Gag was found to recruit G3BP1 and displaces elongation factor 2 or initiation factor 4E-binding protein (4EBP1) in order to block SG assembly^28^, thus counteracting G3BP1-mediated processes, which were reported to sequester HIV-1 RNA transcripts and restrict viral production^29,30^.

In this study, we compared the mRNA and miRNA profiles of uninfected and HIV-1 infected fetal microglial cells. Through gene ontology profiling analyses and by coupling upregulated miRNAs with differentially expressed gene (DEG) mRNAs, we identified and validated miRNA-23b as an HIV-1-induced miRNA that targets the SG core factor gene, *g3bp2* and downregulates its expression. Using monocyte-derived microglial cells (MDMi) as a model, we determined that addition of miRNA-23b mimics reduced the levels of *g3bp2* mRNA and protein, a condition that prevented the formation of SGs in As-treated cells. Consistent with previous reports^28,30^, very few SG were observed in HIV-1 infected cells even under As treatment. Induction of miRNA-23b by HIV-1 was dependent on Env expression and did not involve engagement with CD4 or CCR5. MiRNA-23b inhibitors (antagomirs) impacted negatively the replication and spread of HIV-1 in infected MDMi cultures and partially recovered the formation of SGs in infected cells, a condition phenocopied in MDMi expressing Env-defective HIV-1. Overall, these results underscore the importance of SG assembly in limiting HIV-1 replication and highlight an additional novel mechanism enabling HIV-1 to counter SG assembly during infection of microglia.

## Results

### Transcriptomics of HIV-1 infected microglial cells

Microglial cells isolated from fetal brain samples over 3-weeks of culture were tested for the expression of typical microglia marker proteins (suppl. figure 1A). As expected, cells expressed strong levels of CD11b and c, P2RY12, IBA-1 and TMEM119 ^31^, but also displayed detectable levels of CD4 receptor and CCR5 co-receptor expression, indicating that these cells could be susceptible to HIV-1 infection. The microglial cells were then infected with NL4.3ADA-GFP, a fully-replicative R5-M-tropic HIV-1 modified to additionally express EGFP, as described in Star Methods. Infected GFP-positive (GFP+) and bystander negative (GFP-) cells were sorted following 2 and 7 days of culture (suppl figure 1B), and their mRNA profiles compared to that of uninfected mock (MK) cells (volcano plots, figure 1A). Differentially expressed genes (DEGs) were set at a fold change (fc) > 2 with statistical adjusted *P* values < 0.01. Altogether, 1449 (GFP−/MK, 2 days post-infection (dpi)), 2346 (GFP−/MK, 7 dpi), 1774 (GFP+/MK, 2 dpi) and 2962 (GFP+/MK, 7 dpi) DEGs were identified and analysed. The number of DEGs that were modulated following 2 dpi (D2) exclusively in GFP+ (668 mRNAs) or GFP-(444 mRNAs) subpopulations increased to 1170 mRNAs (GFP+) and 895 mRNAs (GFP-), respectively, after 7 dpi (D7) (figure 1B), suggesting that an important reprogrammation of the transcriptomic profile in both infected GFP+ and GFP-bystander cells takes place over time.

**Figure 1.**
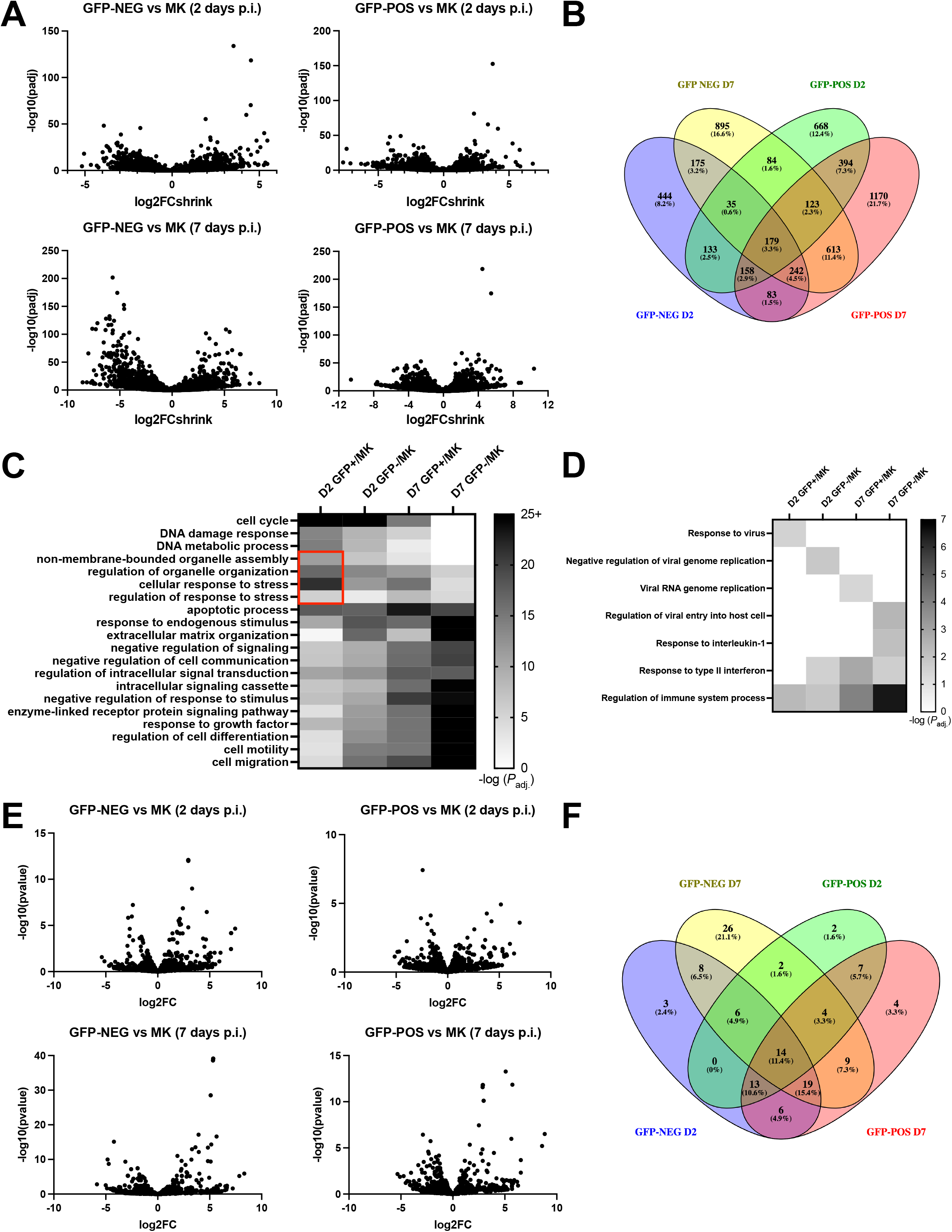
Transcriptomics of HIV-1 infected fetal microglia. A) Volcano plots representing differential expression (fc= “fold change”; padj= “adjusted P value”) of mRNAs in 2-days post-infected (p.i.) or 7 days p.i. GFP+ (GFP-POS) or GFP-(GFP-NEG) fetal microglia *versus* mock infected (MK), derived from 2 donors. B) Venn diagrams showing the distribution of DEGs among GFP+/GFP-cells. D2: 2 days p.i.; D7: 7 days p.i. C) Heatmap of selected gene ontology (GO) biological process (BP) pathways of modulated mRNAs in D2 or D7 GFP+ or GFP-microglial cells, related to mock (MK) infected cells, by −log (adjusted *P*). Pathways related to cellular stress/organelle assembly in D2 GFP+ cells are highlighted in a red square. D) Heatmap of selected GO BP pathways related to immune regulation or virus replication, modulated in cells described as in C. E) MiRNA expression in HIV-1 infected fetal microglia by miRNAseq. Volcano plots representing differential expression (fc= “fold change”; padj= “adjusted P value”) of miRNAs in D2 or D7 GFP+ or GFP-fetal microglia *versus* mock infected (MK), derived from 2 donors. F) Venn diagrams showing the distribution of differentially expressed miRNAs among GFP+/GFP-cells.

To gain a better perspective of the cellular functions altered by the modulation of DEGs in GFP+ or GFP-cells following 2 or 7 days of HIV-1 infection, gene ontology (GO) biological pathways (BP) were analyzed (figure 1C). Several pathways involved in cell motility, cellular differentiation, and intracellular signaling were strongly modulated at D7, particularly in GFP-cells as compared to their D2 counterparts likely reflecting the increased exposure of bystander cells to infected cells; in contrast, BPs relating to cell cycle or DNA damage response/metabolism featured more prominently at D2 in GFP+ and GFP-subpopulations consistent with microglia ability to undergo cell division ^32^. Interestingly, apoptotic processes were most significant in D7 GFP+ infected cells, possibly due to the impact of viral replication over time. BPs related to viral replication and immune response were also modulated although less prominently (figure 1D): for instance, modulated BPs included negative regulation of viral genome replication (*P* adj. ≅ 10^−1.82^; in D2 GFP-cells), response to virus (*P* adj. ≅ 10^−1.49^; in D2 GFP+ cells), as well as regulation of immune system processes in all sorted populations, the latter being more pronounced at D7 (figure 1D). These results suggest that upon infection, microglia undergo profound and progressive transcriptomic changes in genes associated with their activation and capacity to promote immune responses in both the infected and bystander cell populations.

Among the most significantly enriched pathways in D2 GFP+ cells were those related to cellular stress responses (*P* adj. ≅ 10^−21.5^) and non-membrane-bounded organelle assembly (*P* adj. ≅ 10^−11^). Because genes implicated in the formation and regulation of SGs contribute to both pathways and SGs have antiviral functions, we focused subsequent analyses on the associated DEGs.

### Differential expression of miRNAs in HIV-1 infected microglial cells

Similar to a previously described strategy in which we compared both mRNA and miRNA expression in the same sorted cell total RNA samples ^20^, we determined the miRNA expression profile of the uninfected and GFP−/GFP+ microglial cell populations (figure 1E) in order to further gain insights into the functional modulation of cellular genes during HIV-1 infection of microglial cells. Of the total 1541 miRNAs screened (figure 2A), we retained 123 miRNAs (mean read counts ≥ 250; absolute (log (fold change)) ≥ 0.6) for further analyses (figure 1F, suppl. figure 2). Although 26 of the 123 miRNAs were distinctively modulated in D7 GFP-microglia, most of the 123 miRNAs were modulated at both infection time-points or in both GFP−/GFP+ cells, suggesting a broad effect of HIV-1 on miRNA expression in microglial cells (figure 1F). Given the pronounced enrichment of stress response and non-membrane bounded organelle assembly (GO:0140694) BPs in D2 GFP+ cells and the potential indirect effects at D7 due to released soluble factors, we further focused on the miRNAs in D2 GFP+ cells; we also opted to analyse the upregulated miRNAs, given that enhanced miRNA expression may also identify new restrictive processes. For instance, among these upregulated miRNAs were miRNAs-181a/d, which all target SAMHD1, a HIV-1 host restriction factor^33^ (suppl. figures 2 and 3A). We cross-referenced these upregulated miRNAs (suppl. figure 3A) with the genes associated to the GO:0140694 pathway (suppl. figure 3B) to screen for potentially targeted mRNAs using the miRDB database (https://mirdb.org). Only target mRNAs with an miRDB MirTarget^34^ score above 95 were retained.

**Figure 2.**
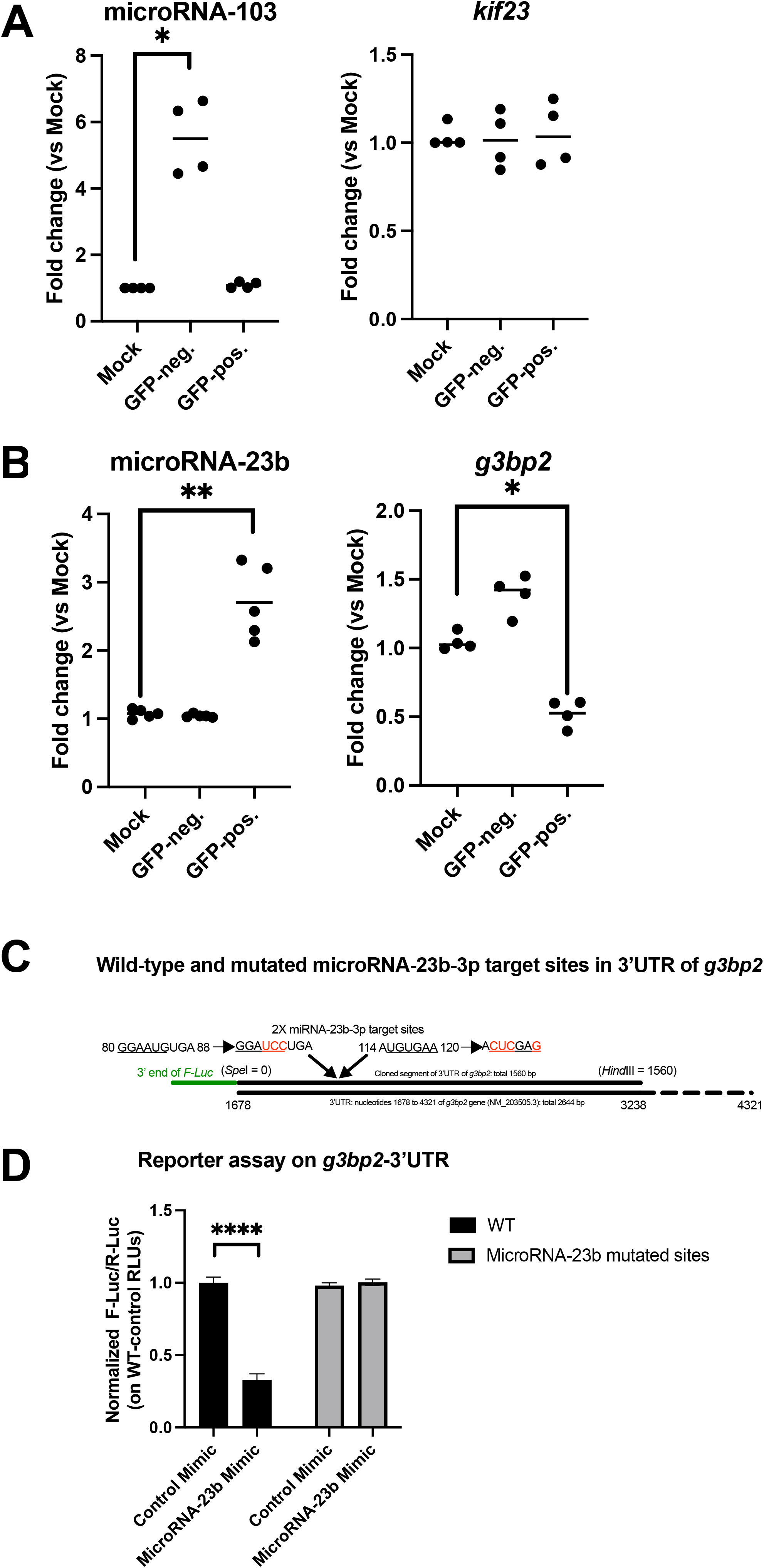
*g3bp2* is a target of miRNA-23b. A) qPCR of miRNA-103 or *kif23* in total RNAs of 2 days infected GFP+ fetal microglial cells (n= 4 donors) (Mann-Whitney test: *, *P* < 0.05). B) Validation of the differential expression of miRNA-23b and *g3bp2* in GFP+ or GFP-fetal microglia (vs mock) by qPCR (n= 4-5 donors) (Mann-Whitney test: *, *P* < 0.05; **, *P* < 0.01). C) The miR-Report validation of the 3’ untranslated region (UTR) of *g3bp2* as a target of miRNA-23b. The WT or mutated (substituted nucleotides in red) versions of the *g3bp2* 3’UTR are shown. The two predicted miRNA-23b target sites are underlined. D) The reporter assay was performed in HEK293T cells transfected with control or miRNA-23b mimics, the Firefly *Luciferase* (F-Luc) reporter plasmids and a Renilla *Luciferase* (R-*Luc*) normalizing control. Relative light units (RLUs) of F-Luc/R-Luc were measured in cell lysates and normalized to control mimics (n=3). Shown are means ±SD of technical duplicates of 3 biological replicates (Student’s *t* test: ****, *P* < 0.0001).

### MiRNA-23b targets the 3’UTR of *g3bp2*

Two miRNA/mRNA couples that displayed an inverse correlation of expression in the GFP+ subpopulation at D2 (suppl. figure 3 A and B) were obtained in the cross-reference screen: *kif23*/miRNA-103a and *g3bp2*/miRNA-23b (suppl. figure 3C). Interestingly, the miRNAseq data (suppl. figure 2) showed that miRNA-23b was also downregulated in both D2 and D7 GFP-cells, indicating that this miRNA is specifically upregulated in HIV-1 infected microglia at D2, as compared to uninfected cells; this was not the case for miRNA-103a, which was broadly upregulated in both GFP+ and GFP-cells at both infection time-points (suppl. figure 2).

We validated the RNAseq/miRNAseq gene expression data on *g3bp2*/miRNA-23b and *kif23*/miRNA-103a in fetal microglial cells by quantitative real-time PCR (qPCR). Microglial cells obtained from different donors were infected with NL4.3ADA-GFP for 2 days and sorted for GFP-expression. We were unable to confirm either *kif23* or miRNA-103a as DEGs in D2 HIV-1 infected GFP+ microglial cells by qPCR (figure 2A). However, the downregulation of *g3bp2* and increase of miRNA-23b at D2 in the GFP+ sorted cells versus uninfected cells were observed in samples from all donors (figure 2B), supporting the data obtained by RNAseq/miRNAseq. This led us to focus our investigation on *g3bp2* and miRNA-23b.

To verify that *g3bp2* mRNA is a target of miRNA-23b, the first 2/3 of the 3’UTR of the *g3bp2* gene, which contains two potential miRNA-23b binding sites (figure 2C and suppl. figure 3C), was cloned and fused to the Firefly *Luciferase* (*F-Luc*) reporter gene. The addition of miRNA-23b mimics into HEK293T cells expressing the chimeric *F-Luc*-*g3bp2*-3’UTR reporter plasmid resulted in a decrease of F-Luciferase activity in cell lysates as compared to control mimics-transfected cells (figure 2D); this was not the case for cells expressing a chimeric *F-Luc-g3bp2*-3’UTR in which both miRNA-23b target sequences in the *g3bp2*-3’UTR were mutated (figure 2C). Taken together, these results validate *g3bp2* as a target of miRNA-23b and further suggest that an increase of miRNA-23b early post infection (D2) leads to a decrease in *g3bp2* mRNA in HIV-1 infected primary microglial cells.

### HIV-1 glycoproteins expression drives miRNA-23b induction, which is conserved among several HIV-1 strains in infected monocyte-derived microglia (MDMi)

In order to further investigate the potential mechanisms involved in miRNA-23b-mediated *g3bp2* mRNA downregulation and its consequences in HIV-1 infection, we proceeded to use monocyte-derived microglial cells (MDMi) ^35,36^ as a microglia model, which circumvented the limitations associated with access to primary microglial cells from multiple donors. When fully differentiated from human monocytes, MDMi expressed microglia markers P2RY12, TMEM119 and IBA-1 as well as the HIV-1 receptor CD4 and coreceptor CCR5 (figure 3A). Importantly, upon D2 infection with NL4.3ADA-GFP, sorted GFP+ MDMi showed reduced expression of *g3bp2* mRNA and increased miRNA-23b expression (figure 3B), similarly to that observed in primary microglial cells (figure 2). Furthermore, transfection of miRNA-23b mimics into MDMi cells significantly reduced *g3bp2* mRNA as compared to control mimic-transfected cells (figure 3C); this was not the case for mRNA of the closely related *g3bp1* gene (figure 3C). Taken together, these results indicate that the interplay between miRNA-23b and its specific target *g3bp2* can be recapitulated in the MDMi model of microglia upon infection with HIV-1. To verify that the miRNA-23b/*g3bp2* mRNA modulation in D2 HIV-1 infected GFP+ MDMi is not limited to the NL4.3ADA strain, four other HIV-1 constructs (also expressing GFP) were tested and analysed: the macrophage-tropic AD8 ^37^, the neurotropic YU2c ^20^ and primary HIV-1 transmitted/founder isolates THROc ^38^ and WITO ^39^. GFP+ MDMi, infected 2-days (indicated in green) with either AD8, YU2c, THROc or WITO GFP-expressing strains showed a similar pattern of expression of *g3bp2* and miRNA-23b as with NL4.3ADA-GFP (figure 3D), indicating that the modulations of *g3bp2* and miRNA-23b is a conserved property of other R5-M-tropic HIV-1 strains. As previously reported ^10^, infection of MDMi with the CXCR4 (X4)-tropic NL4.3-GFP viruses did not result in productive infection (no GFP+ cells, indicated in black) and the recovered virus exposed cells did not exhibit any enhancement of miRNA-23b or reduction of *g3bp2* mRNA expression (figure 3E). However, NL4.3-GFP viruses that were pseudotyped with VSV-G did infect MDMi (indicated in green) and did upregulate miRNA-23b expression in GFP+ cells to a degree comparable to NL4.3ADA-GFP, suggesting that X4 viruses have the ability to upregulate miRNA-23b as long as viral entry or viral expression in MDMi is achieved (figure 3E). Although pseudotyping Env-deficient NL4.3-GFP with either HIV-1 ADA Env or VSV-G restored productive infection (indicated in green), neither condition increased miRNA-23b nor altered *g3bp2* expression in GFP+ MDMi (Figure 3E). Overall, these results suggest that receptor/co-receptor engagement does not mediate the increase in miRNA-23b and point to a role of *de novo* HIV-1 glycoproteins expression in triggering miRNA-23b induction in infected MDMi.

**Figure 3.**
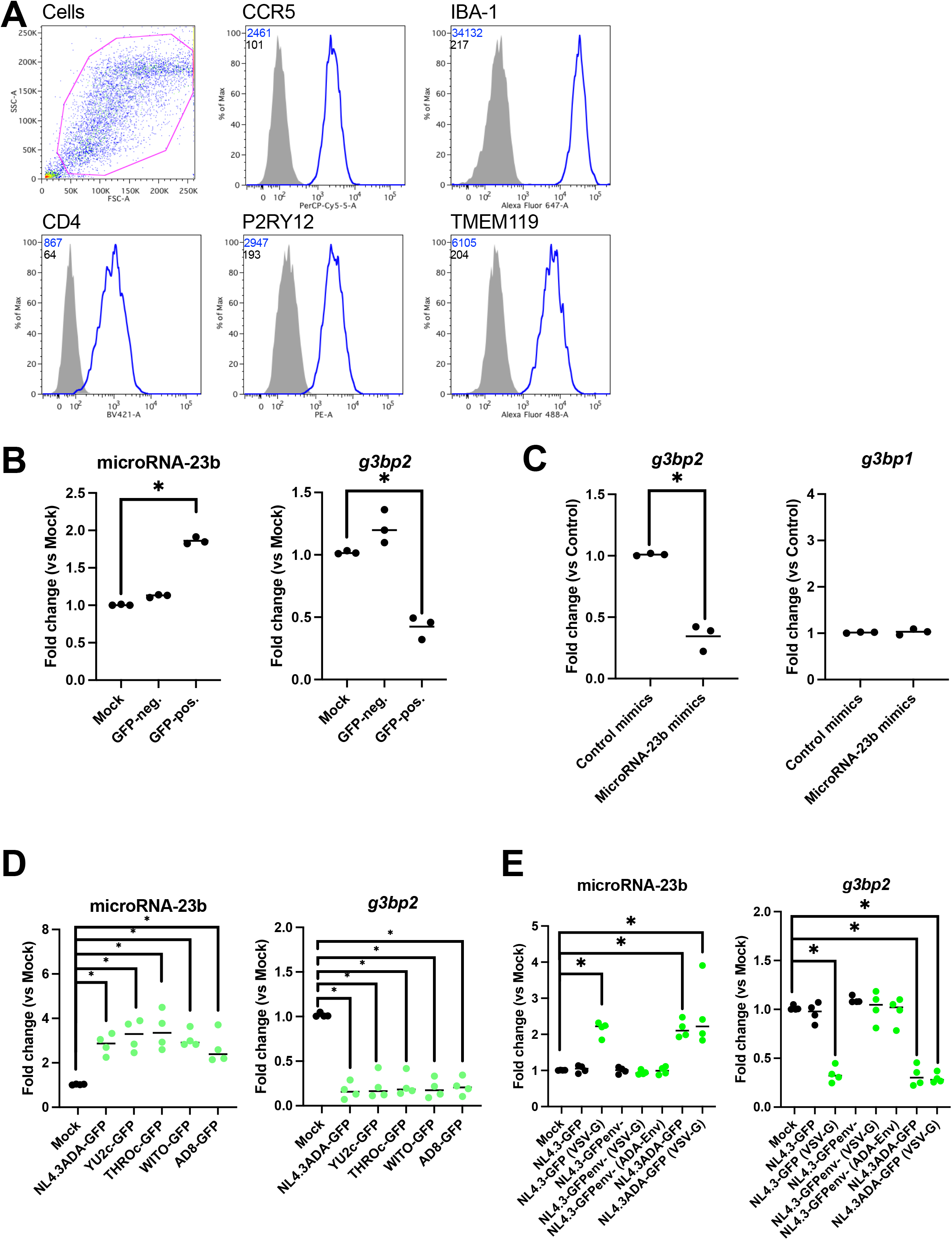
Monocyte-derived microglial cells (MDMi) as a model of HIV-1 infection of microglia. A. Characterization of MDMi. Cells were stained for the indicated surface receptors or intracellular protein (IBA-1), fixed and analyzed by flow cytometry. MFI obtained with the control isotypic (dark) or specific antibodies (blue) are shown in a representative example. B. The expression of miRNA-23b and *g3bp2* in HIV-1 infected GFP+ or GFP-MDMi (vs mock) at D2 was determined by qPCR (n= 3 donors). C. The expression levels of *g3bp2* and *g3bp1* mRNAs were quantified by qPCR in control or miRNA-23b-mimic transfected MDMi (n= 3 donors). D. Expression profile of miRNA-23b and *g3bp2* is similar in MDMi infected with the different indicated HIV-1 strains. Green dots represent conditions of productive infection and gene expression in sorted GFP+ MDMi (*vs* mock-infected, in black) infected for 2 days with HIV-1 GFP-marked HIV-1 strains; expression levels of miRNA-23b and *g3bp2* were determined by qPCR (n= 4 donors). All viruses were pseudotyped with VSV-G to optimize the infection rate. E. HIV-1-mediated miRNA-23b upregulation is dependent on Env glycoproteins expression. Green dots represent conditions of productive infection and gene expression in sorted GFP+ MDMi (*vs* mock-infected, in black) infected for 2 days with the indicated HIV-1 GFP-marked constructs; expression levels of miRNA-23b and *g3bp2* were determined by qPCR (n= 4 donors). Black dots represent conditions where the indicated virus did not lead to productive infection. For consistency, cells under these conditions were ‘mock’ sorted. Viral glycoproteins used to pseudotype virus particles are shown in parentheses (Mann-Whitney test: *, P < 0.05).

### MiRNA-23b modulates stress granule formation

The G3BP2 protein and its paralog G3BP1 are essential constituents of SG assembly^40,41^ and function as nucleators of SGs to potentiate their formation following most stress conditions, for instance, those induced by the addition of As ^22^. Given the effect of miRNA-23b on *g3bp2* mRNA, we determined by confocal microscopy imaging if the addition of miRNA-23b mimics interfered with the formation of SGs in MDMi induced by As treatment. MDMi exhibited a typical round or ovoid body with primary processes and secondary branches^10^ (figure 4A). Although SGs-containing G3BP1 and G3BP2 were visible in control mimic-treated MDMi following the addition of As, a marked reduction in the frequency of cells displaying G3BP2-positive SG as well as in the numbers of G3BP2-positive SG per cell was observed in miRNA-23b mimic-treated cells (figure 4A-B). These data indicate that miRNA-23b, by modulating G3BP2 expression, can regulate SG formation in these cells, even though the levels of *g3bp1* mRNA remain unaffected in the presence of miRNA-23b mimics (figure 3C). The reduction of G3BP2 protein levels (figure 4C-D) in MDMi treated with miRNA-23b mimics was further validated by western blotting of cell lysates. MiRNA-23b mimics reduced G3BP2 levels without altering G3BP1 expression in both As-treated and untreated cells (figure 4C-D).

**Figure 4.**
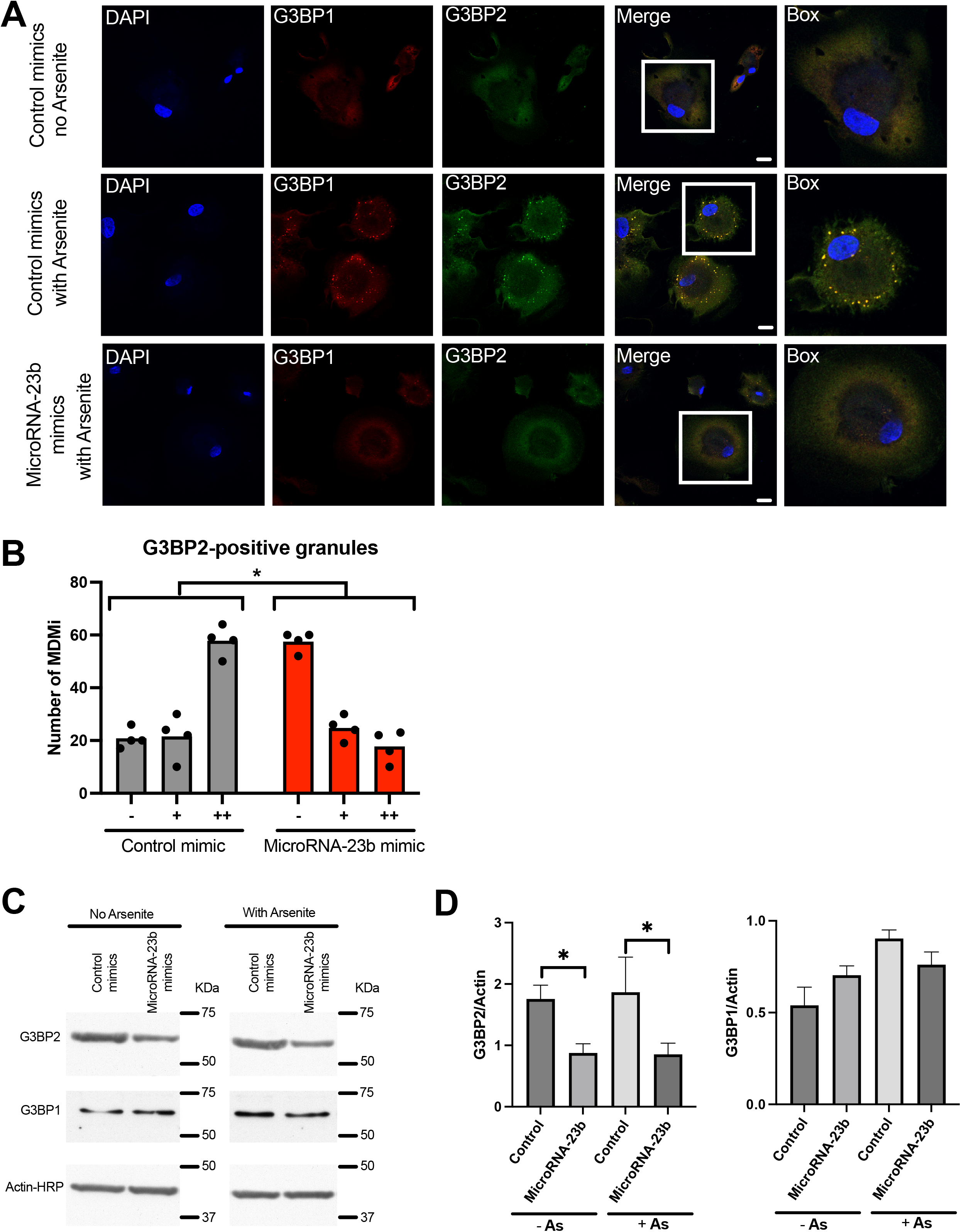
Assembly of SGs and levels of G3BP2 protein are reduced in miRNA-23b mimic-transfected MDMi under conditions of cellular stress. A. MDMi were transfected with control or miRNA-23b mimics, treated or not with As, stained for G3BP1 (red) and G3BP2 (green) and imaged using confocal microscopy to visualize SGs. Bar = 5 μm. B. Quantitation of cells containing G3BP2-positive SGs as observed in A. Symbols “-”, “+”and “++” represent 0, 1-5 and >5 SGs/cell, respectively (Kolmogorov-Smirnov test: *, P < 0.05; n=100 cells/condition, n= 4 donors, histograms represent the means). C. Control or miRNA-23b mimics transfected MDMi were untreated or treated with As, lysed and the levels of G3BP1 and G3BP2 proteins determined by Western immunoblotting. Quantitation of protein levels as determined by densitometry is shown in D (means ±SD, of n=4, biological replicates) (Mann-Whitney test: *, P < 0.05).

### MiRNA-23b inhibitors reduce HIV-1 replication and spread in MDMi

Given that HIV-1 upregulates miRNA-23b expression in infected microglial cells and that this miRNA reduces SG formation, we set out to determine what was the impact of miRNA-23b on HIV-1 replication and spread in MDMi derived from three distinct donors. MDMi were treated with either miRNA-23b mimics or inhibitors (antagomirs) prior to infection and cultured for two weeks post-infection; viral spread and replication were measured by the percentage of GFP-positive cells over time and HIV-1 production assessed by the level of HIV-1 p24 in the cell supernatants, respectively. As previously reported, infection of MDMi by R5-M-tropic HIV-1 led to efficient virus production and spread^10^ (figure 5). MiRNA-23b antagomir-treated infected cultures showed a greatly reduced HIV-1 spread and production, particularly starting at day 6 post-infection (figure 5). Comparatively, miRNA-23b mimics induced a detectable increase in HIV-1 spread and production in MDMi cultures generated from all three donors (figure 5). These data indicate that miRNA-23b positively regulates HIV-1 infection in MDMi, likely by interfering with the restrictive activity exerted by G3BP2-mediated SG formation.

**Figure 5.**
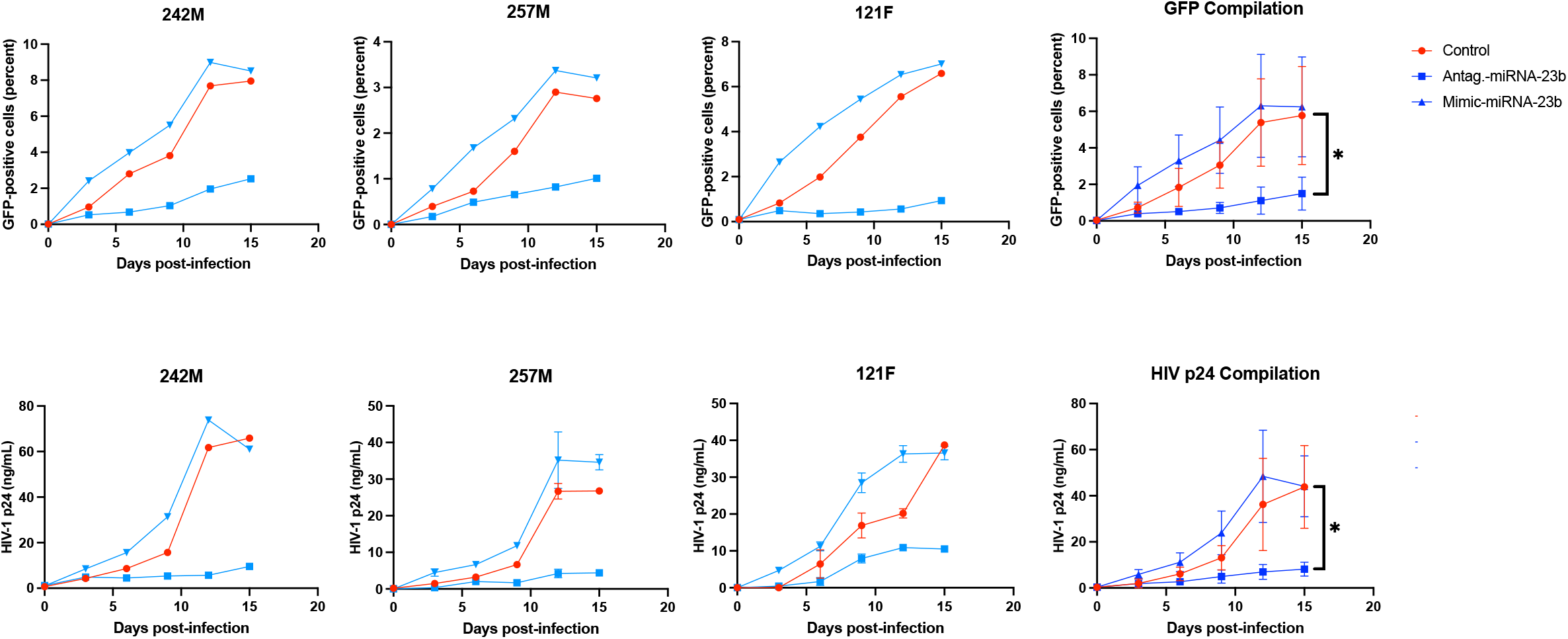
Effect of miRNA-23b antagomirs or mimics on HIV-1 replication and spread in MDMi. Control, miRNA-23b mimics or antagomir-transfected MDMi were infected with NL4.3ADA-GFP and cultured for 15 days. A. The spread of infection was determined by evaluating the percentage of GFP+ cells in the culture at different time intervals. B. Virus production was determined at different time points by assessing the levels of HIV-1-associated p24 in the culture supernatant. Compilation of data from MDMi derived from the 3 different blood donors are shown (Wilcoxon rank test: *, P < 0.05).

### Interfering with miRNA-23b partially restores SG formation in HIV-1 infected MDMi

HIV-1 Gag has been reported to counter SG assembly by targeting G3BP1 in virus-expressing cells, and the lack of SG formation counteraction was found to result in impaired HIV-1 production and infectivity^28^. To determine if the increase of miRNA-23b in HIV-1 infected cells contributes to the counteraction of the restrictive G3BP2-mediated SG formation, in addition to that exerted by Gag, confocal microscopy analyses of 2-day HIV-1 infected GFP+ MDMi stained for several SG components, including G3BP2 and G3BP1, were conducted (figure 6). Rare G3BP1- or G3BP2-containing SGs were detected in infected GFP+ MDMi cell cultures, consistent with induction of miRNA-23b during infection (figure 3B) and the reported countermeasure elicited by HIV-1 Gag on SG assembly ^28^ (figure 6A). The quantitation of SGs in GFP+ MDMi was indeed very similar to that of bystander GFP-MDMi (figure 6B). We next examined the impact of miRNA-23b inhibitors (antagomirs) on SG formation. As shown in figure 6A, treatment with miRNA-23b antagomirs induced a detectable increase (∼ two-fold) in both the frequency of SG-positive cells (containing G3BP1 or G3BP2) and the actual number of SGs per cell in the GFP+ infected MDMi subpopulation. This increase was not observed in the GFP-bystander cells, of which quantitation remained similar to control (figure 6A and B). This was also the case for SGs containing PABPC1, which is also an important factor in SG assembly ^24^ (suppl. figure 4). However, it is noteworthy that the vast majority (>80 %) of infected GFP+ MDMi remained exempt of detectable SGs (figure 6B, figure suppl 4).

**Figure 6.**
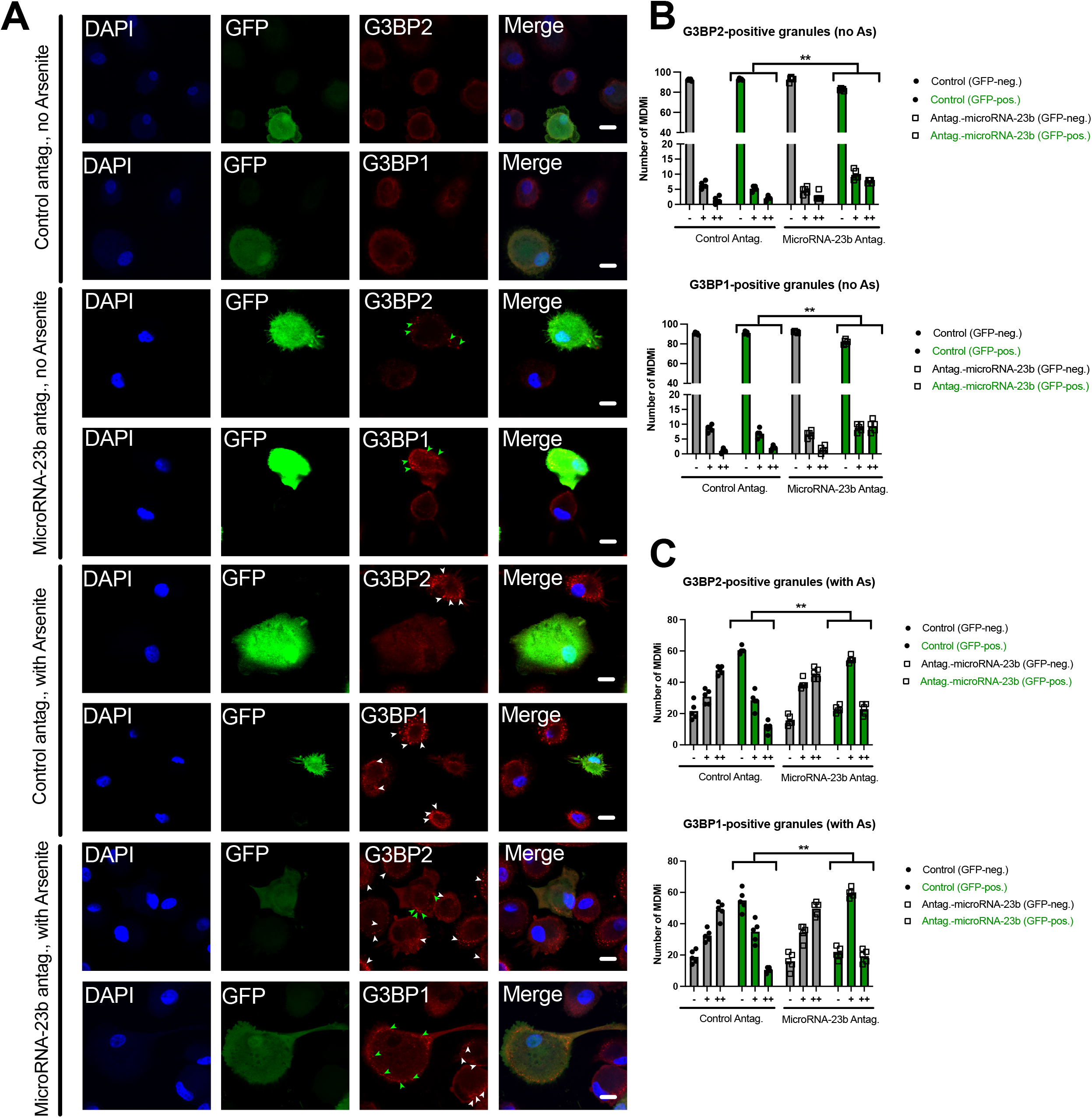
Treatment with miRNA-23b antagomirs promotes the formation of SGs in HIV-1 infected (GFP+) MDMi. A. HIV-1 infected MDMi were transfected with control or miRNA-23b antagomirs and treated or not with As, as indicated. Cells were fixed, stained for either G3BP1 or G3BP2 and imaged using confocal microscopy. Arrows point to SGs in infected GFP+ (green) or non-infected bystander GFP-(white) cells. Bar = 5 μm. B and C. Quantitation of MDMi, GFP-(grey) or GFP+ (green) containing 0, 1-5 and >5 SGs/cell, as represented by symbols “-”, “+”and “++”, respectively, in As-untreated (B) and As-treated (C) cell cultures, as observed in A. Histograms represent the means (Kolmogorov-Smirnov test: *, P < 0.05; n=100 cells/condition; n=5 donors).

In order to confirm the differences in SG assembly observed between infected GFP+ and bystander GFP-MDMi in the presence of miRNA-23b antagomirs, a similar set of analyses were performed in HIV-1 infected MDMi treated with As (figure 6A and C). The addition of As resulted in GFP-cells assembling far more G3BP1/2 SGs than their GFP+ counterparts, which remained mostly exempt of SG (figure 6C, control antagomir-treated cells), suggesting that, in this context, HIV-1 mediated countermeasures that interfere with SG formation are more apparent. This was also the case for SGs containing PABPC1 (suppl. figure 4). Interestingly, treatment with miRNA-23b antagomirs allowed infected GFP+ MDMi to recover their ability to assemble As-induced SGs (both in terms of frequency of cells harboring SGs (about two-fold) and number of SGs per cell) (figure 6A and C, suppl. figure 4), although not to the levels of their GFP-counterparts, of which numbers remained similar to the control GFP-MDMi (figure 6A and C, suppl. figure 4).

Having previously shown that HIV-1 Env glycoproteins are a determinant in the induction of miRNA-23b in infected MDMi (figure 3E), we tested for the impact of VSV-G pseudotyped, Env-negative HIV-1 on the formation of SGs in cells treated or not with As. As shown in figure 7, infection with VSV-G pseudotyped Env-negative viruses partially recovered formation of SGs (identified by G3BP1 or G3BP2 staining) in GFP+ MDMi, as compared to fully replicative WT viruses; this was the case in both As-treated or untreated cells (figure 7).

**Figure 7:**
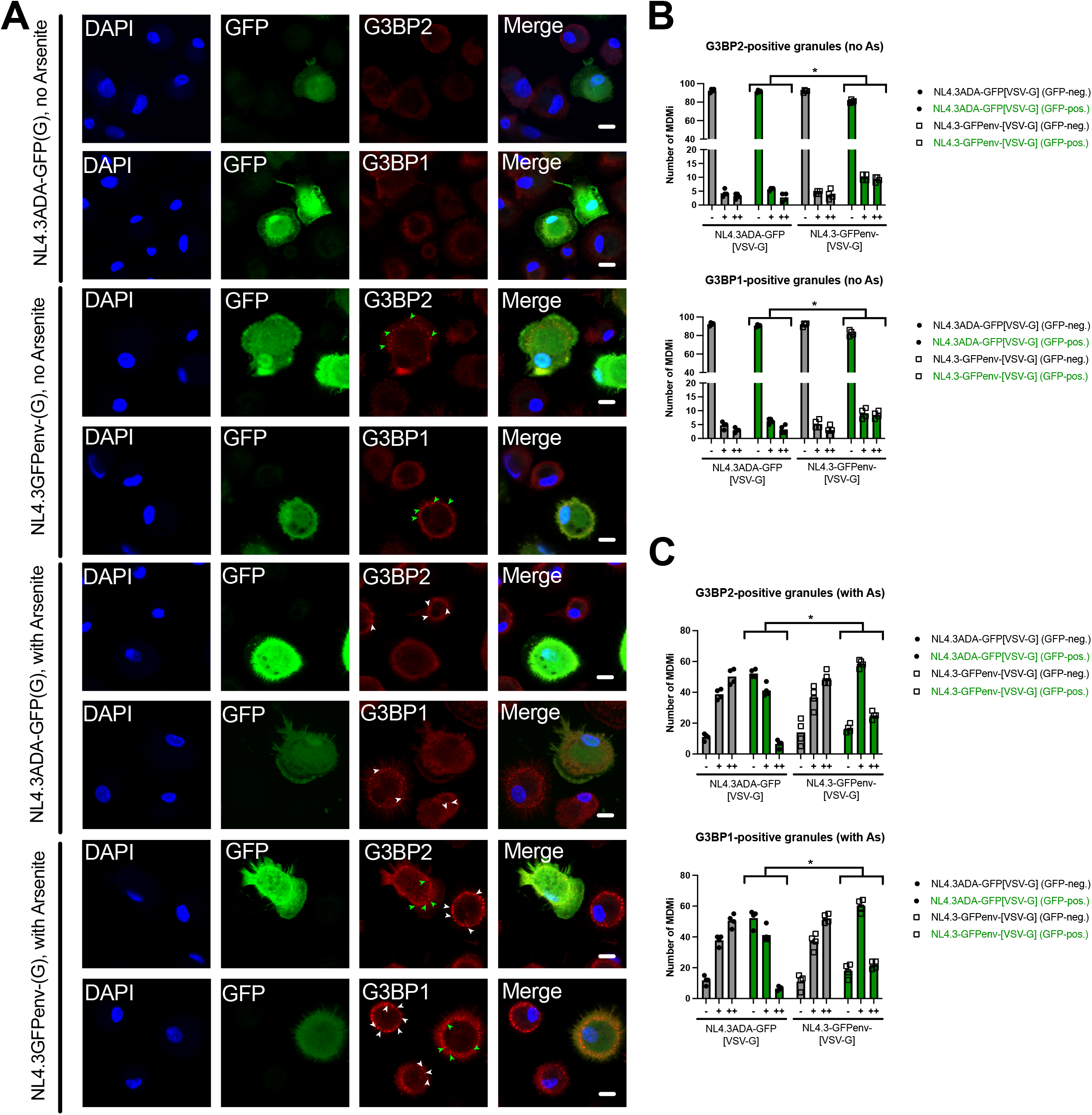
Formation of G3BP1/2-containing SGs in MDMi infected with Env-negative HIV-1 pseudotyped with VSV-G. A. MDMi were infected with WT or Env-negative (both VSV-G pseudotyped) HIV-1 particles and treated or not with As, as indicated. Cells were fixed, stained for G3BP1 or G3BP2 and imaged using confocal microscopy. Arrows point to SGs in infected GFP+ (green) or non-infected bystander GFP-(white) cells. Bar = 5 μm. B and C. Quantitation of MDMi GFP-(grey) or GFP+ (green) containing 0, 1-5 and >5 SGs/cell, as represented by symbols “-”, “+”and “++”, respectively, in As-untreated (B) and As-treated (C) cell cultures, as observed in A. Histograms represent the means (Kolmogorov-Smirnov: *, P < 0.05; n=100 cells/condition; n=4 donors).

Taken together, these data support the notion that Env-mediated induction of G3BP2-targeting miRNA-23b expression during HIV-1 infection of microglia contributes to inhibition of SG assembly, a condition that promotes HIV-1 replication and spread. The ability to reduce viral replication and spread despite a partial recovery of SG formation, when miRNA-23b activity is inhibited or its upregulation prevented, suggests that these biocondensates exert a potent restriction on HIV-1 replication beyond the actual infected cells where SG formation is restored.

## Discussion

Our gene ontology analyses of the biological pathways from transcriptomic mRNA profiles of productively HIV-1 infected (GFP+) and bystander (GFP-) fetal microglial cells revealed an enhanced modulation of cell cycle and DNA damage/metabolic pathways in D2 GFP+ cells and to a lesser degree in D7 GFP+ cells. Among the genes whose expression was upregulated was *p21* (*CDKN1A*), a hallmark of p53 activation (figure 1). Interestingly, cell cycle/DNA damage/metabolic pathways modulation was also detected in D2 GFP-cells, which correspond to latently infected or uninfected bystander microglia, but this enrichment could not be detected by D7 (figure 1C). Our transcriptomics data analyses identified also modulated pathways involved in both viral replication and immune/cytokine regulation (figure 1D); the changes in immune processes were more enhanced through time (D2 to D7), in line with the concurrent spread of HIV-1 infection.

Gene ontology analyses determined also that the pathways related to the assembly of non-membrane-bounded organelles and cellular response to stress were more prevalent in GFP+ D2 infected cells (figure 1C). Indeed, several HIV-1 proteins have been shown to induce oxidative stress, including Vpr ^48^, Tat ^49^ and gp120 ^50^. Overall, our transcriptomics data of modulated pathways in productively HIV-1 infected (GFP+) and bystander (GFP-) fetal microglial cells is reminiscent of the findings of recent studies using single-cell RNA sequencing of HIV-1 infected human induced pluripotent stem cells (iPSC) derived microglia ^47^ and profiling of biomarkers in infected primary fetal microglia ^42^, which provided evidence that HIV-1 infection induced a pattern of cellular senescence in microglia. This senescence signature was characterized by marked upregulation of genes promoting cell-cycle arrest, the downregulation of proliferation-associated genes, and extensive remodeling of metabolic-stress and inflammatory-signaling pathways in both infected and bystander cells. Given that dysregulated microglial function contributes to HAND pathogenesis, these findings provide a molecular framework for how HIV-1 infection may accelerate neurocognitive decline in infected individuals^43^.

MicroRNA profiling of infected microglia at D2 and D7 identified 123 miRNAs that were differentially expressed in the infected GFP+ and bystander GFP-subpopulations relative to the mock control (figure 1 and figure suppl 2). While previous studies reported the induction by the HIV-1 transactivator Tat of several miRNAs, including miRNA-32 ^44^, miRNA-34a ^45^ and miRNA-505 ^46^ involved in the activation of microglia, none of these miRNAs were found to be upregulated in our miRNA profiling of infected GFP+ or bystander GFP-primary microglia (suppl figure 2). The fact that these miRNAs and their targets were uncovered in experimental cellular models upon exposure to recombinant HIV-1 Tat protein might explain their absence in our miRNA profiling studies.

To identify new candidate genes involved in regulating HIV-1 infection of macrophages, a previous approach was used that paired both miRNAseq and RNAseq data^20^. Analyses of modulated mRNAs of genes associated to non-membrane-bounded organelle assembly coupled to upregulated miRNAs (suppl. figure 3) identified miRNA-23b and potential target *g3bp2* as possible host factors involved in modulating HIV-1 infection of microglia. The induction of miRNA-23b and the concomitant downregulation of *g3bp2* expression during early infection of microglia was found to be a feature shared by several isolates of HIV-1 (figures 2, 3), suggesting that this conserved activity is biologically relevant.

MiRNA-23b has been reported to target *g3bp2* in kidney tissues and the HK-2 kidney epithelial cell line ^47^, and to be downregulated by p53 ^47,48^. MiRNA-23b is also downregulated by the pro-inflammatory cytokine IL-17, which initiates the binding of IκB-ζ to NF-κB sites in the miRNA promoter ^49^. Among the anti-inflammatory roles of miRNA-23b is its targeting of *TAB2* and *TAB3*, that are involved in TNF-α/IL-1β-mediated NF-κB activation ^49^. Particularly relevant, miRNA-23b has been associated to several neuroinflammatory diseases, such as Alzheimer’s disease ^50^. Given its anti-inflammatory functions, miRNA-23b upregulation in HIV-1 infected microglia would be a mechanism that counters the activation state of these cells and perpetuates the viral reservoir. Interestingly, *de novo* synthesis of the HIV-1 glycoproteins, but not engagement of the CD4 receptor and CCR5 co-receptors, is necessary for increasing levels of miRNA-23b in HIV-1 infected microglial cells; this could be shown by using Env-negative viruses (figure 3E) While the mechanism of induction of miRNA-23b involving HIV-1 Env remains unclear, the miRNA-inducing ability of Env is not unique to miRNA-23b. Indeed, it was recently reported by our group that HIV-2 Env is involved in triggering the expression of miRNA-25 and miRNA-93 to counter the expression of MARCH1, a protein which restricts HIV infectivity by limiting Env incorporation into nascent virions ^20,51^.

G3BP1 and 2 both participate in SG formation and are among the core initiators of their assembly ^23^; both proteins also reinforce the integrated stress response translation programme ^52^. Besides their role in SG formation, G3BP1/2 have other functions, such as tethering the Tuberous sclerosis complex (TSC) to lysosomes to suppress Master regulator mechanistic target of rapamycin complex 1 (mTORC1) signaling ^53^, leading to cell migration and metabolic disorders. It is noteworthy that although G3BP2 is about 4-times more expressed than G3BP1 in the brain, both proteins are similarly expressed in microglial cells ^54^. MiRNA-23b mimics-treatment of MDMi reduced the number of As-induction of SGs (figure 4A, B) and the levels of G3BP2 protein (figure 4C) although G3BP1 mRNA (figure 3C) and protein (figure 4C) levels remained unchanged. This suggests that G3BP2 is necessary for efficient As-induced SG assembly in MDMi, even in the presence of G3BP1, consistent with the fact that these two core components are essential for initiating SG assembly ^23^.

SGs and other membraneless organelles, such as RNAse-L induced bodies^55^, cGAS foci^56^ and protein kinase R clusters^57^, act in antiviral defense mechanisms following the interaction of host pattern recognition receptors with viral-derived ligands^58^. The antiviral activity of SGs mostly resides in their sequestering of untranslated viral RNAs, disrupting viral replication cycles^59^. Given that SGs function as antiviral platforms ^60^, viruses have evolved strategies to counter this function. For example, G3BP1 mediates restriction of HIV-1 infection by sequestering viral RNA in T-cells and macrophages ^29^; however, HIV-1 Gag recruits G3BP1 which leads to SG dissassembly^28^ favoring HIV-1 production and replication. Our results support the notion that the induction of miRNA-23b in HIV-1 infected MDMi to counter G3BP2-mediated SG assembly is yet an additional viral strategy to limit SG formation and their ensuing restriction of HIV-1 infection; indeed, inhibiting miRNA-23b using specific antagomirs greatly reduced viral production and spread in infected MDMi cultures (figure 5). Furthermore, when treated with miRNA-23b antagomirs, infected GFP+ MDMi partially recovered their ability to assemble SGs that contained G3BP1/2 and PABPC1, an observation confirmed in conditions of stress induced by As (figure 6, suppl. figure 4). Thus, either inhibiting miRNA-23b (antagomirs, figure 6) or affecting its induction by preventing HIV-1 Env expression leads to a reduction in the formation of SGs (figure 7). An incomplete recovery is to be expected given that SG components, such as G3BP1 ^28^, are also targeted by HIV-1 to counter SG formation. While these results establish a link between the ability of miRNA-23b to modulate G3BP2-mediated SG formation and the degree of viral replication and spread (figures 5, 6), the extent of the inhibition of viral production and spread in miR-23b antagomir-treated cells (figure 5), even in a context of a partial recovery of SGs (figure 6), suggests that a limited recovery of SG formation is sufficient to exert a potent restrictive effect on viral infection; alternatively, we cannot exclude that miRNA-23b may have another/other target(s), in addition to *g3bp2*, that limit HIV-1 replication in infected microglia.

Taken together, these findings uncover a new mechanism that is significant for our understanding of how viruses can counter antiviral stress responses and notably the assembly of SGs.

### Limitations

Our study has limitations that should be acknowledged. First, given the difficulty to obtain samples to isolate primary fetal microglia, transcriptomics data were obtained from only 2 donors, albeit both samples showed consistent clustering for HIV-infected (GFP+/GFP-) and uninfected subsets on PCA scatter plots. Second, primary fetal microglia may differ from adult CNS microglia, although both of these cultured cellular models display similar morphologies and immune functions, including phagocytosis and inflammatory responses, comparable expression of microglia core genes and HIV-relevant cellular genes and both models are susceptible to CCR5-tropic HIV-1 infection^10^. Finally, although sorted HIV-1 infected primary fetal microglial cells were used to first identify *g3bp2* and miRNA-23b, the effect of miRNA-23b on SGs was mechanistically shown using MDMi. MDMi are easier to obtain and offer technical advantages over primary fetal microglia, but it is noteworthy that microglia are not totally recapitulated in the MDMi monocyte-derived model ^10,61^. In this regard, it would be interesting to further investigate our findings using microglia grown in 3D cerebral organoids^10^ or *in vivo* models, which better recapitulate the complex cellular interactions and regulatory mechanisms of the brain environment. Nevertheless, both miRNA-23b and *g3bp2* were modulated accordingly in primary fetal microglial cells and MDMi.

## Resource availability

### Lead contact

Requests for further information and resources should be directed to the lead contact, Eric A. Cohen.

### Materials availability

Material generated in this study will be shared by the lead contact upon request.

### Data and code availability

Data reported in this paper is available upon request to the lead contact.

The online GEO accession numbers of the mRNAseq and miRNAseq data reported in this study are GSE351612 (mRNAseq) and GSE351611 (miRNAseq), respectively. The mRNAseq data is derived from two samples, and the miRNAseq data is derived from the same samples, corresponding to donors 1 and 2 (of 4) presented in the miRNAseq.

Any additional information required to reanalyze data reported in this paper is available from the lead contact upon request.

## Acknowledgements

We thank Tina Lee (IRCM Clinic) for coordinating access to blood donors, as well as the participants. We thank the IRCM and the CR-CHUM Flow Cytometry Cores for their assistance with cell sorting, and the IRCM Molecular Biology and Bioinformatics Cores for the processing and analyses of the RNAseq/miRNAseq samples and data, respectively. We thank Dr. Florence Margottin-Goguet for the pSIV3+ plasmid. The following reagent was obtained from the NIH AIDS Reagent Program: TZMbl cells ARP-8129 from Dr. John C. Kappes, Dr. Xiaoyun Wu, and Tranzyme Inc. We are grateful to Grégoire de Bisschop, Nicolas Jacques-de Vries and Éric Lécuyer (IRCM) for their gifts of antibodies and helpful discussions. We thank Thomas Durcan (McGill University Montreal Neurological Institute) and members of the Canadian HIV-Ageing Multidisciplinary Programmatic Strategy (CHAMPS) in NeuroHIV Research Team for helpful discussions.

This study was supported by the Canadian Institutes of Health Research (CIHR) grants FDN-154324 to É.A.C. and HB2-164064 (Canadian HIV Cure Enterprise—CanCURE) to É.A.C. and C.P.; É.A.C. is the recipient of the IRCM-Université de Montréal Chair of Excellence in HIV Research. FK was supported by the Deutsche Forschungsgemeinschaft (DFG, Ki548/21-1; SFB 1506, C01) and an ERC Advanced grant (Traitor viruses).

## Author contributions

Conceptualization: RL, EAC; methodology/reagents: RL, WB, FK, CP, EAC; experimentation: RL; Data analyses: RL, EAC; writing-original draft: RL, EAC; writing-review and editing: RL, WB, FK, CP, EAC; funding acquisition: EAC, CP; supervision: EAC, CP.

## Star Methods

### Experimental model and study participants details

#### Primary cell samples and participants

Human primary microglia cultures were prepared from fetal tissue obtained from 15-20 weeks electively terminated healthy pregnancies with written informed consent of the donors (Pro000027660), as approved by the University of Alberta Human Research Ethics Board (Biomedical) and the IRCM Human Research Ethics Review Board (IRCM 2019-954).. Peripheral blood samples were obtained from HIV- and HCV-seronegative healthy adults (of either gender) who gave written informed consent in accordance with the Declaration of Helsinki under research protocols approved by the IRCM Human Research Ethics Review Board (IRCM 2012-16). The number of blood donors from which cells were used is mentioned in figure legends.

#### Primary cell culture

Human fetal microglia were prepared as previously reported ^62^. Briefly, following dissection of fetal brain tissue, dissociated cells were obtained by digesting with 0.25% trypsin (Lifetech) and 0.2 mg/ml DNAse I (Roche) and passage through a 70 μm cell strainer. Cells were washed twice in culture medium and plated in flasks. Cells were maintained in MEM supplemented with 10% fetal bovine serum (FBS), 2 mM L-glutamine, 1 mM sodium pyruvate, 1X MEM non-essential amino acids, 0.1% dextrose, 100 U/ml Penicillin, 100 μg/ml Streptomycin, 20 μg/ml gentamycin and 0.5 μg amphotericin B (Fungizone, Gibco). After 2 weeks of cell culture, loosely attached or free-floating microglia were recovered by gentle rocking of the plate, separating them from more adherent astrocytes and neurons, washed and plated in 6 wells plates. Characterization of microglia was assessed by flow cytometry analyses. Monocyte-derived microglial (MDMi) cells were derived from human peripheral blood mononuclear cells (PBMCs). Briefly, PBMCs from healthy blood donors were generated from buffy coats following Ficoll density gradient separation (GE Healthcare) as previously described ^19^. MDMi were obtained, as previously described ^36^ by differentiating plate-adhered monocytes from PBMCs in RPMI-1640 supplemented with 10% FBS, L-glutamine, 100 ng/ml IL-34 (Biolegend) and 10 ng/ml GM-CSF (Peprotech) for 14 days. Differentiated MDMi were characterized by flow cytometry.

#### Cell lines

HEK293T cells (ATCC) were generated from epithelial cells that were originally isolated from a female fetal kidney tissue. The TZM-bl cell line^63^ was acquired from the NIH AIDS reagent program; it was generated from HeLa cells, a cell line derived from a 30 year old female with papillomavirus-related endocervical adenocarcinoma. HEK293T and TZM-bl cells were maintained in DMEM media with 10% FBS and Penicillin-Streptomycin.

### Method details

#### Chemicals and antibodies

Chemicals and antibodies used were: Live-Dead Fixable yellow (Molecular Probes, Invitrogen, ThermoFisher); APC mouse anti-CD11b ICRF44, PerCP-Cy5.5 mouse anti-CD11c 3.9, PerCP-Cy5.5 rat anti-CCR5 J418F1, PerCP-Cy5.5 mouse anti-CD4 OKT4, PE mouse anti-P2RY12 S16001E, mouse anti-TMEM119 A16075D from Biolegend; mouse anti-IBA-1 MABN92, mouse anti-G3BP1 O5-1938, rabbit anti-PABPC1 ABE-40 from Millipore-Sigma; rabbit anti-G3BP2 A302-040A (Bethyl), mouse anti-beta-actin-HRP AC15 (Abcam) and Sodium arsenite (NaAsO2, Honeywell Fluka 35000-1L). The mimics of miRNA-23b-3p (339173-YM00472291-ADA) and miRCURY LNA antagomirs of miRNA-23b-3p (339121-YIO4102299-ADA) were from Qiagen (see Table 1).

#### Flow cytometry

The expression of a range of microglia markers was determined by flow cytometry. Cells were recovered by gentle scraping after a 10 minute, 37 °C incubation in PBS-EDTA (5 mM), washed in FACS buffer (PBS containing 2% FBS) and Fc receptors blocked with human IgGs (Sigma) in FACS buffer (blocking buffer). For intracellular marker IBA-1, 0.1% Triton X-100 was added to blocking buffer for 15 minutes to permeabilize cells. Cells were then stained with indicated fluorochromed-labeled antibodies, or unlabeled antibodies, followed by extensive washes in PBS. Unlabeled antibodies were further stained with the indicated AlexaFluor-labeled antibodies. Following washes, cells were fixed in 4% paraformaldehyde in PBS, resuspended in PBS-EDTA, and analyzed on a BD Biosciences Fortessa cytometer equipped with appropriate lasers. Detailed analyses were obtained using the FlowJo software package (version 9.3.2) and geometric mean fluorescence intensities were recorded.

#### G3BP2 3’UTR validation assay

The miR-REPORT system (Applied Biosystems, Ambion) was used to validate *g3bp2* as a miRNA-23b target, with plasmid pGL4.70Actin1.2-8 encoding the *Renilla* luciferase replacing pMiR-REPORT-βgal. The pMiR-REPORT-G3BP2(H3)-Luc was generated by first performing *Pfu* PCR on cDNA of primary peripheral blood monocytes using primers g3bp2-utr-WT-Fwd and g3bp2-utr-WT-hindIII (H3)-Rev, which produces a fragment encompassing about 2/3 of the 2644 bp *g3bp2* 3’ UTR (GenBank: NM_203505.3) up to a unique HinDIII restriction site at position 3238; this 1560 bp fragment was then inserted into the pMiR-REPORT-Luc SpeI and HinDIII restriction sites. The pMiR-Report-G3BP2(H3)-Luc plasmid harboring nucleotide substitutions in the two sequences recognized by miRNA-23b in the *g3bp2* 3’UTR was obtained by site-directed mutagenesis overlap PCR, using primers MiR-23b-siteA-Fwd and MiR-23b-siteA-Rev (for the first target sequence), MiR-23b-siteB-Fwd and MiR-23b-siteB-Rev (for the second target sequence) in combination with g3bp2-utr-WT-Fwd or g3bp2-utr-WT-hindIII-Rev, as appropriate (see Table 1). The 3’UTR validation was performed in HEK293T cells transfected with the pMIR-REPORT-G3BP2(H3)-Firefly-Luc-derived plasmids, pGL4.70Actin1.2-8 encoding the Renilla-Luc and control or miRNA-23b mimics using Lipofectamine RNAi-max (ThermoFisher Scientific) and read-outs were assessed using the Dual-Glo luciferase assay system (Promega) on a GloMax luminometer (Promega).

#### Plasmids, viruses and lentiviral vectors

HIV-1 proviral DNA plasmids NL4.3-ADA-GFP-IRES-NEF (described as “NL4.3ADA-GFP” in the text)^64^, NL4.3-GFP-IRES-NEF ^65^, and Env-negative NL4.3ΔEnv-GFP-IRES-NEF ^66^ were described elsewhere. Briefly, NL4.3-ADA-GFP-IRES-NEF was generated by transferring the 2.7 kb SalI-BamHI fragment of NLHXADA into the NL4.3 backbone at these two sites. The SalI-BamHI cassette contains a SacI-BsmI insert encoding for ADA Vpu and Env gp120 and two flanking sequences (SalI-SacI and BsmI-BamHI) of HXB2 which encode for the first and second exons of Tat and Rev. The insertion of this DNA fragment rendered NL4.3 macrophage-tropic ^67^ through the ability of its recombinant Env containing the ADA determinant to mediate viral entry via a CCR5-dependent mechanism ^19^. HIV-1 YU2c, THRO and AD8 proviral reporter constructs containing a functional *nef* gene followed by an IRES and the *eGFP* gene were generated as described for NL4.3^68^. WITO-GFP-IRES-NEF was created by inserting the GFP-IRES fragment from NL4.3-ADA-GFP-IRES-NEF into WITO ^38,39^ by PCR using primers WITOEndEnv-NotI-GFP FWD and IRES-WITO Nef REV (Table 1). This GFP-IRES fragment was cloned into the BamHI and XhoI sites of WITO by overlap PCR using GFP-IRES in combination of PCR products PCR-A (WITOBamHI FWD and WITOEndEnv REV) and PCR-B (WITONef FWD and WITOXhoI REV) (Table 1). The viral glycoprotein-expressing plasmids SV-CMV-VSV-G ^69^ and pSV-III-Env-ADA ^70^ were also previously described. The SIV3+ plasmid ^71^ was used to produce Vpx-containing lentiviral particles. Virus stocks were produced in HEK293T cells by calcium phosphate transfection, virus titers calculated using TZMbl cells ^63^ and concentration of HIV p24 determined by ELISA p24 (Xpress Bio) as previously described ^19^.

#### HIV-1 infection, mimic/antagomir transfection

Primary fetal microglia and MDMi were pre-treated with Vpx-containing SIV3+ particles prior to HIV-1 infection, in order to counter any SAMHD1 remaining restriction^71^. MDMi were mock-infected or infected with VSV-G pseudotyped HIV-1 unless otherwise indicated at a MOI of 1 and retrieved by gentle scraping. Cells were sorted on a BD FACSAria Fusion (BD Biosciences) flow cytometer for GFP expression (infected GFP+ and bystander GFP-) and directly recovered in RLT lysis buffer (Qiagen) for RNA extraction and kept at −80 °C. In some cases, MDMi were transfected with control or miRNA-23b mimics or antagomirs using Lipofectamine RNAi-max (ThermoFisher Scientific), and the percentage of GFP-expressing infected MDMi determined on a BD Biosciences Fortessa cytometer following cell fixation in PBS with 4% paraformaldehyde. Virus production was measured by the levels of HIV-1 p24 in the cell supernatant by p24-ELISA (Xpress Bio). Primary microglia were mock-infected or infected with NL4.3-ADA-GFP-IRES-NEF at an MOI of 1 for either 2 or 7 days and sorted for GFP expression using an Influx (BD Biosciences) cell sorter. Sorted microglia were directly recovered in RLT lysis buffer (Qiagen) and kept at −80 °C.

#### RNA extraction and qPCR

RNeasy RNA columns (Qiagen) were used to extract total cellular RNAs (including miRNAs) according to the manufacturer’s instructions, and samples stored at −80 °C. Reverse transcription of RNAs was performed using SuperScript III or IV reverse transcriptase with poly(dT) using primers described in Table 1; two-tailed qPCR ^72^ loop primers designed for specific miRNAs (Table 1) were added to the mix. For qPCR, cDNAs and appropriate primers (Table 1) were added to SYBR green select master mix (Applied Biosystems) in 96-well plates and run on a ViiA96 thermocycler (ThermoFisher Scientific). *GAPDH* was used as the loading control. ΔΔ*CT* variations determined and calculated relative to those of uninfected cells.

#### SGs in mimic-transfected MDMi

MDMi were transfected with control or mimics for miRNA-23b for 24 hours, washed and treated with 500 µM As in PBS for 30 minutes. Cells were then either processed for microscopy or lysed and processed as described previously for Western immunoblotting ^73^. Protein lysates were separated by sodium dodecyl sulfate-polyacrylamide gel electrophoresis, and western immunoblotting performed as previously described ^73^.

#### Microscopy of SGs in MDMi

MDMi were seeded on coverslides, infected for 2 days with NL43ADA-GFP-IRES-NEF viruses, and treated with control or antagomirs for miRNA-23b for 2 days. Cells were then washed, treated with 500 µM As (or not) in cell culture media for 30 minutes, washed and fixed in 4% w/v paraformaldehyde in PBS. Cells were then kept in PBS, permeabilized in 0.1% Triton X-100 and Fc receptors blocked with human IgGs (Sigma). Cells were then stained with antibodies against SG-containing proteins, washed, and stained with anti-rabbit and/or anti-mouse fluorescent antibodies combined with DAPI. Following final washes, cells were mounted on slides and imaged with a Zeiss 700 or 710 confocal microscope. Images were analysed with the NIH Image software package.

#### RNAseq of microglia

Differential gene expression in HIV-1 infected or uninfected control fetal microglia was compared using the Illumina system (Illumina Technologies) at the IRCM Molecular Biology and Functional Genomics Core Facility. Specific tagging was used to identify RNA from each sample originating from 2 donors. The resulting libraries were sequenced at the Core Facility; 50 bp paired-end (PE50) sequencing was used on a HiSeq 2000 sequencer (Illumina Technologies). The quality of the raw reads was assessed with FASTQC v0.12.1. After examining the quality of the raw reads, no trimming was deemed necessary. The reads were aligned to the GRCh38 reference genome with STAR v2.7.11b. The raw counts were calculated with FeatureCounts v2.0.6 based on the GRCh38 reference genome (release 98). Differential gene expression was performed using the DESeq2 R package. Differentially expressed gene (DEG) heatmaps were drawn based on log(Fold change)shrinck. Functional enrichment analyses of DEGs (gene ontology and pathway enrichment) were performed with the gprofiler2 R package using FDR correction. Significant results were reduced using term size between 15 and 2000, and intersection size > 2. The mRNAs with an adjusted *P* value < 0.01 and Fold change > 2 were considered for further analyses. Bioinformatics analyses were performed at the IRCM Bioinformatics Core Facility. Volcano plots were generated using GraphPad Prism9 (GraphPad Software, LLC) and Venn graphs using Venny 2.1 (https://bioinfogp.cnb.csic.es/tools/venny/).

#### MiRNAseq of microglia

Sample RNAs from mock or HIV-1 infected fetal microglia originating from 2 donors were processed using the Illumina TruSeq Small RNA system, and libraries sequenced as above. Adapter sequences were trimmed using cutadapt, and reads were filtered to 22 nucleotides to retain mature miRNA species. Processed reads were mapped to the reference genome using miRDeep2 (‘mapper.pl‘), followed by miRNA quantification with ‘miRDeep2.pl’ against miRbase v22. Differential expression analyses were performed using DESeq2 in R. Counts were normalized using the median-of-ratios method, and statistical significance was assessed using Wald tests with Benjamini–Hochberg correction. Only miRNAs with a base mean above 250 and absolute log(Fold change) > 0.6 were considered for further analyses. Identification of miRNA sites on 3’UTRs was obtained using Targetscan (Human, v.8.0). The online search for miRNA and mRNA pairing was performed using mirRDB (https://mirdb.org).

### Quantification and statistical analysis

#### Statistics

Statistical analyses were performed on GraphPad Prism 9 using the non-parametric Mann-Whitney rank test, Student’s t test, or one-way ANOVA. SG distribution statistics on microscopy data were performed using the Kolmogorov-Smirnov test in GraphPad Prism 9. Statistical significance is indicated in the figures: *, *P* < 0.05; **, *P* < 0.01; ***, *P* < 0.001; ****, *P* < 0.0001. Statistics for *P* values for the RNAseq and miRNAseq differential expression data were calculated using DESeq2.

