## Supplemental_files_and_Table for "Impact of HIV-1 infection on the microglia transcriptome: induction of microRNA-23b leads to G3BP2-mediated stress granules assembly disruption"

Figure suppl 1

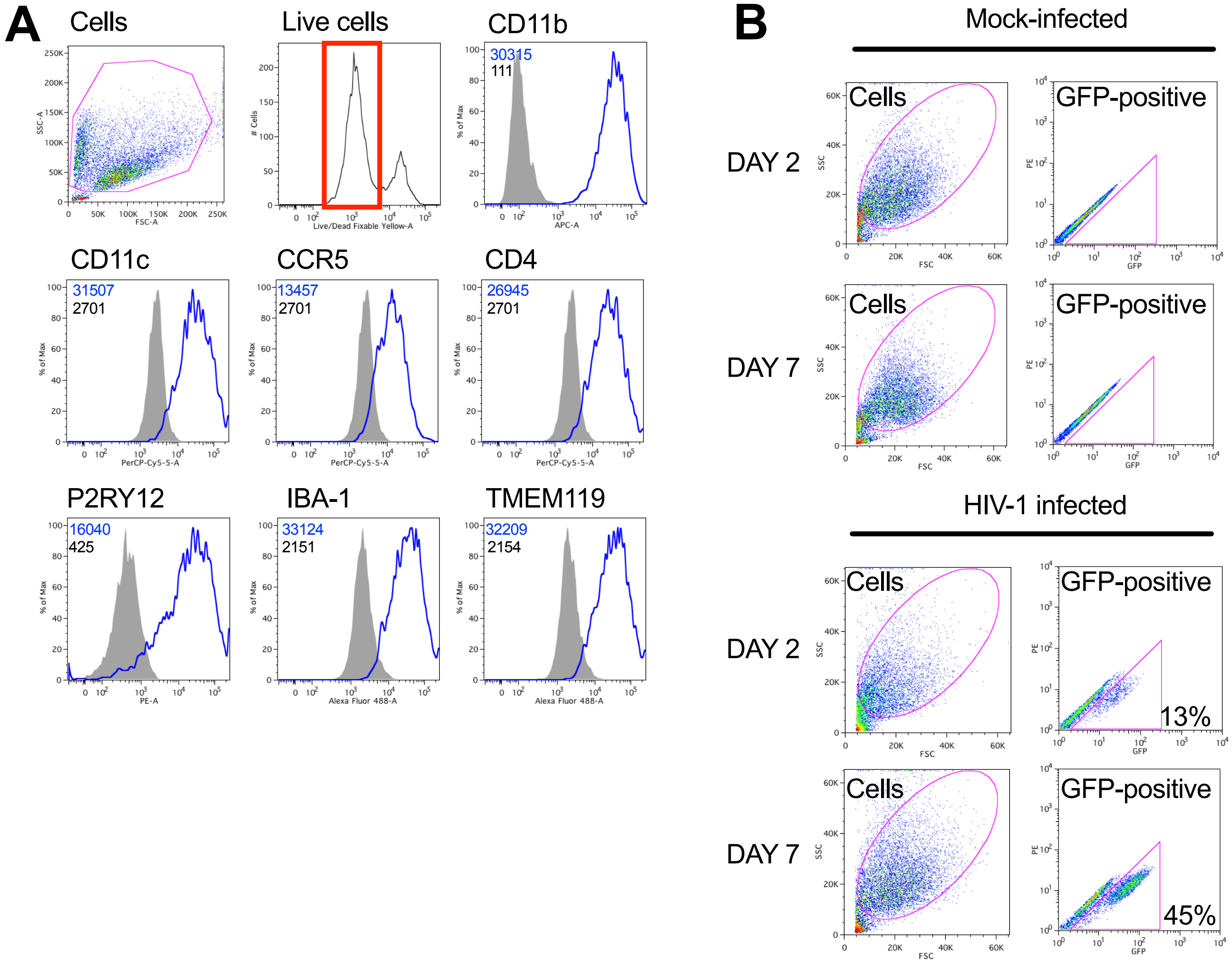


***Supplementary figure 1**: **Characterization, infection and sorting of fetal microglial cells.** A. Microglial cells were stained for the indicated receptors or intracellular protein (IBA-1) , fixed and analyzed by flow cytometry. Mean geometric fluorescent intensities (MFI) obtained with the isotypic controls (dark) or with the indicated specific antibodies (blue) are shown in a representative example. B. Representative example of the GFP+/GFP- sorting of 2 or 7 days infected microglial cells (percentages of infected GFP+ cells are indicated). Mock-infected microglial cells after 2 and 7 days of culture are shown as control. The increased frequency of GFP+ infected cells over time is indicative of a spreading infection.*

Figure suppl 2

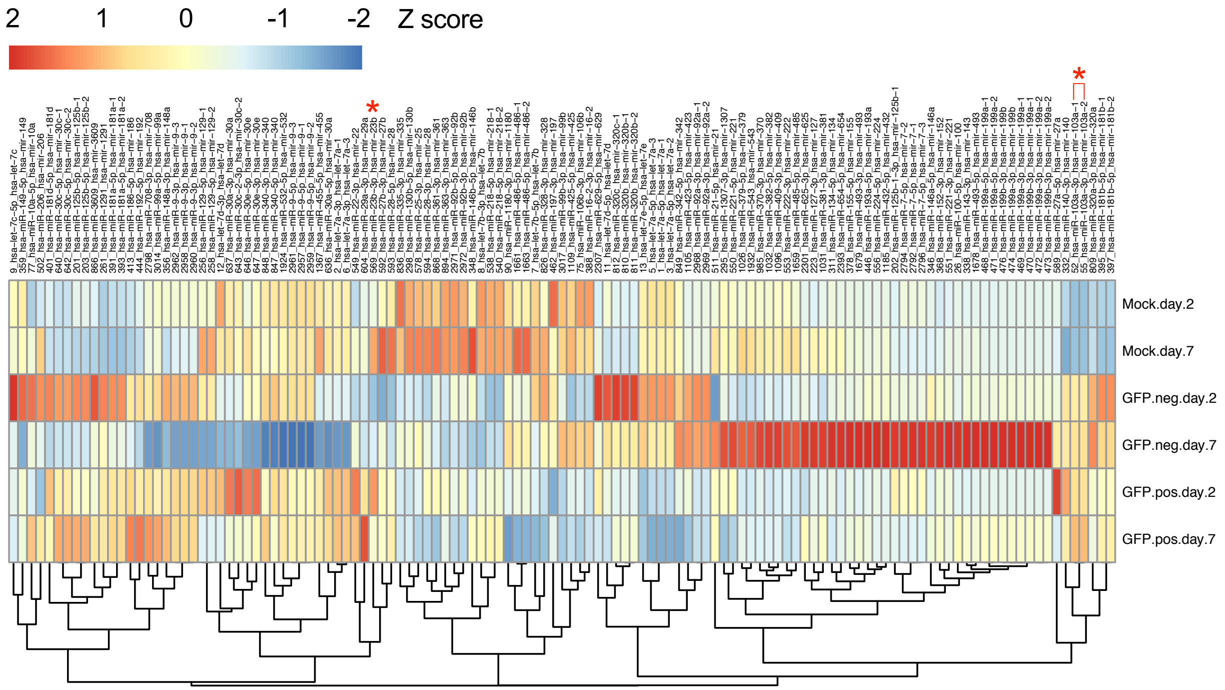


***Supplementary figure 2*: *Heatmap of differentially-expressed miRNAs*.** *Shown are miRNAs that have a base mean > 250 and absolute logFC > 0.6; Z-scores are presented to allow for analyses between the different sorted GFP+/GFP-/mock populations. MicroRNAs of interest are shown with an asterisk (*).*

Figure suppl 3

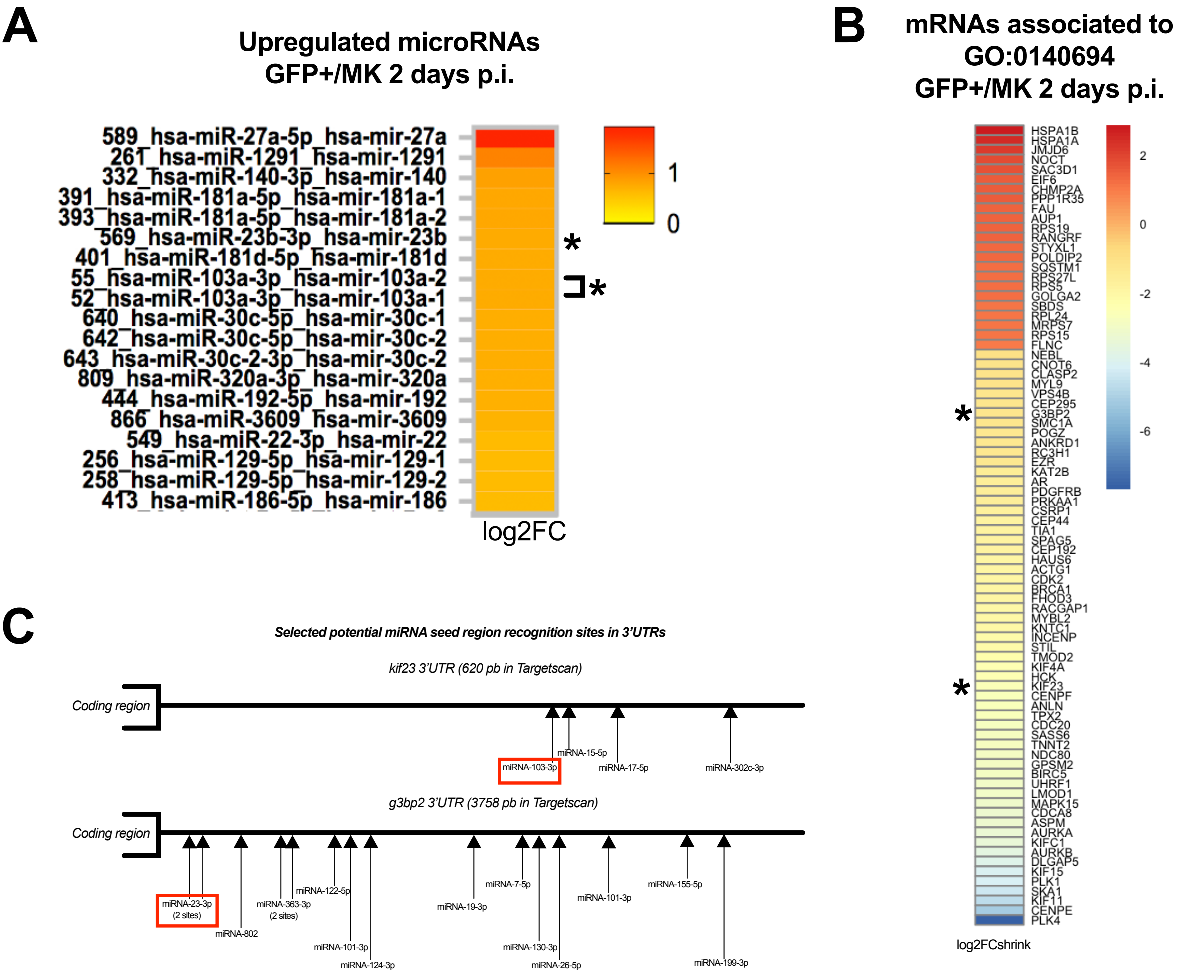


***Supplementary figure 3**: MiRNA/mRNA paired analyses in 2-day GFP+ HIV-infected fetal microglia.* *A. Heatmap of the upregulated miRNAs in the 2 day-infected GFP+ fetal microglial cells, versus MK. Asterisks indicate the miRNA/mRNA pairs according to MiRDB (https://www.mirdb.org) cross-analyses, where only miRNAs with a MiRDB MirTarget score of > 95 for every mRNA were retained. B. Heatmap of modulated mRNAs in the 2 day-infected GFP+ microglial cells, versus uninfected (MK), associated with GO:0140694 (membrane-less organelle assembly). C. Schematic representation of the kif23 and g3bp2 3' untranslated regions (3'UTR) with Targetscan (www.targetscan.org) predicted miRNA target sites shown.*

Figure suppl 4

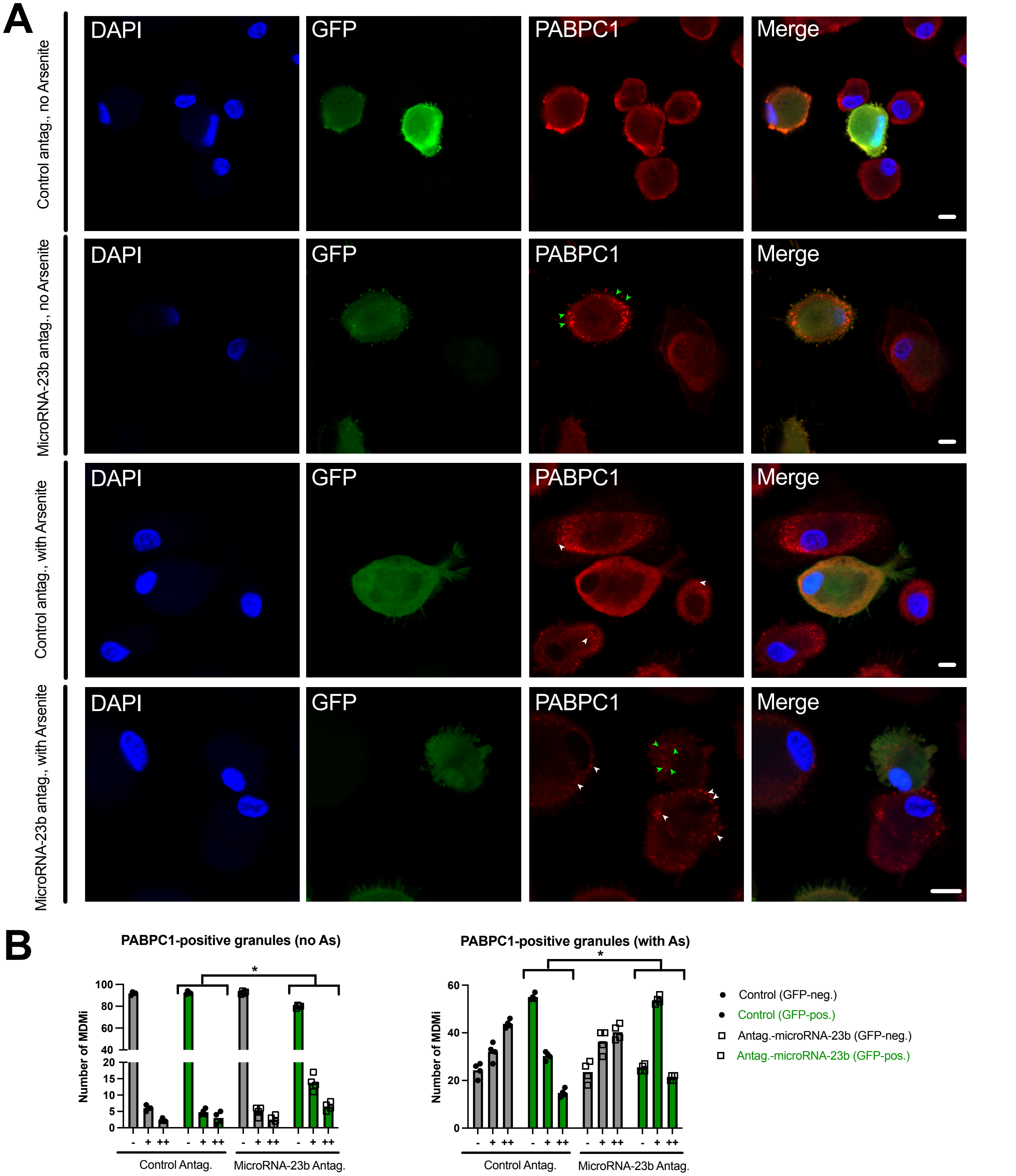


***Supplementary figure 4**: **Treatment with miRNA-23b antagomirs promotes the formation of PABPC1-containing SGs in HIV-1 infected (GFP+) MDMi**. A. HIV-1 infected MDMi were transfected with control or miRNA-23b antagomirs and treated or not with As, as indicated. Cells were fixed, stained for PABPC1 and imaged using confocal microscopy. Arrows point to SGs in infected GFP+ (green) or non-infected bystander GFP- (white) cells. Bar = 5 μm. B. Quantitation of MDMi, GFP- (grey) or GFP+ (green) containing 0, 1-5 and >5 SGs/cell, as represented by symbols "-", "+"and "++", respectively, in As-untreated (left) and As-treated (right) cell cultures, as observed in A. Histograms represent the means. The Kolmogorov-Smirnov test was used for statistical analyses (*, P < 0.05; n=100 cells/condition; 4 donors).*

Table 1: Oligonucleotides used in this paper.

| Oligonucleotide | Sequence |
|---|---|
| **MicroRNA-23b mimics and antagomirs** | |
| MiRNA-23b antagomir | GGTAATCCCTGGCAATGTGA |
| MiRNA-23b mimic | AUCACAUUGCCAGGGAUUACC |
| **Primers for qPCR** | |
| G3BP1-FWD | AAACGTTTGTCCTTGCTCCT |
| G3BP1-REV | TTCAGACTCCTCCTGAGGCT |
| G3BP2-FWD | GAGCTGAAACCACAAGTGGAGG |
| G3BP2-REV | GGTCACTGAAGCCCAGGAGAAA |
| GAPDH-FWD | GCCATCAATGACCCCTTCAT |
| GAPDH-REV | TTGACGGTGCCATGGAATTT |
| KIF23-FWD | TGCCATGAAGTCAGCGAGAG |
| KIF23-REV | GCGCACCCTACAGTATACCC |
| MiRNA-23b-FWD | GCAATGTGGATATGTGAGACGTACGTTG |
| MiRNA-23b-REV | GGATCACATTGCCAGGGATTACCAC |
| MiRNA-23b-Loop | GCAATGTGGATATGTGAGACGTACGTTGAGTACGTCAAGTGAAGTGTGGTAATCC |
| MiRNA-103-FWD | ACAATGTCAAGCTCTCCAGGTACAGT |
| MiRNA-103-REV | GGAGCAGCATTGTACAGGGCT |
| MiRNA-103-Loop | TACAATGTCAAGCTCTCCAGGTACAGTTGGTACCTGACTCCACGCTCATAG |
| **MiR-REPORT constructs** | |
| g3bp2-utr-WT-FWD | ACTAGTAGCTCCACTGTTGGCAAAGTCTTG |
| g3bp2-utr-WT-(H3)-REV | GGATAAGCTTTGCCATACTTGTGGTTTACTG |
| MIR-23b-siteA-FWD | GCTTTGGATCCTGACACAGCCTTTTTG |
| MiR-23b-siteA-REV | CAAAAAGGCTGTGTCAGGATCCAAAGC |
| MiR-23b-siteB-FWD | GATCATTTCTTTGACTCGAGAAGCATCTTTGGTTATC |
| MiR-23b-siteB-REV | GATAACCAAAGATGCTTCTCGAGTCAAAGAAATGATC |
| **WITO-GFP-IRES-NEF** | |
| WITOEndEnv-NotI-**GFP** FWD | GGGCTTCGAAAGGGCTTTACTATAAGATGGGGGGCGCGGCCGCAATGGTGAGCAAGGGCGAGG |
| **IRES**-WITONef REV | CCATCCAATTTTCCAACTTTTTGACCACTTGCCCCCCATATTATCATCGTGTTTTTCAAAGGAAAACCACG |
| WITO**BamHI** FWD | GGAGGGGATCCAGAAGTTGTAATGC |
| WITO**EndEnv** REV | TTATAGTAAAGCCCTTTCGAAGCCCTGTC |
| WITO**Nef** FWD | ATGGGGGGCAAGTGGTCAAAAAG |
| WITO**XhoI** REV | CCATGTCTTTCTAGGTCTCGAGATACTGC |
